# Locally low-rank denoising in transform domains

**DOI:** 10.1101/2023.11.21.568193

**Authors:** Steen Moeller, Erick O. Buko, Suhail P. Parvaze, Logan Dowdle, Kamil Ugurbil, Casey P. Johnson, Mehmet Akcakaya

## Abstract

**Purpose:** To develop an extension to locally low rank (LLR) denoising techniques based on transform domain processing that reduces the number of images required in the MR image series for high-quality denoising.

**Theory and Methods:** LLR methods with random matrix theory-based thresholds are successfully used in the denoising of MR image series in a number of applications. The performance of these methods depend on how well the LLR assumption is satisfied, which deteriorates with few numbers of images, as is commonly encountered in quantitative MRI applications. We propose a transform-domain approach for denoising of MR image series to represent the underlying signal with higher fidelity when using a locally low rank approximation. The efficacy of the method is demonstrated for fully-sampled k-space, undersampled k-space, DICOM images, and complex-valued SENSE-1 images in quantitative MRI applications with as few as 4 images.

**Results:** For both MSK and brain applications, the transform domain denoising preserves local subtle variability, whereas the quantitative maps based on image domain LLR methods tend to be locally more homogeneous.

**Conclusion:** A transform domain extension to LLR denoising produces high quality images and is compatible with both raw k-space data and vendor reconstructed data. This allows for improved imaging and more accurate quantitative analyses and parameters obtained therefrom.

## Introduction

Signal to noise ratio (SNR) in MRI is proportional to the voxel size and the square root of the acquisition time. Thus, with technological advances that strive towards improved spatial resolutions and faster imaging, many scans end up in a depleted SNR regime. The need for higher resolutions and faster imaging is even more pronounced in applications where multiple MR images are acquired across time or different contrasts, including functional MRI (fMRI)[1], MR relaxometry[2], diffusion MRI (dMRI)[3], arterial spin labeling (ASL)[4], and dynamic contrast-enhanced MRI[5].

To improve SNR, numerous techniques have been proposed for denoising of MR image series, including those based on low-rank approaches[6-13], block matching self-similarity[14], and nonlocal means[15]. Low-rank techniques are either applied globally to the whole image series or in a local manner to image patches[12]. The latter technique, referred to as locally low-rank (LLR) regularization, has been widely used for MR image series such as dMRI[7-9, 11], fMRI[10], and ASL[16], while also being used for cardiac imaging[12], multiparametric mapping[17], and relaxometry[18, 19]. More recently, a line of work that utilizes ideas from random matrix theory has established parameter-free denoising for removing signals that cannot be distinguished from independent and identically distributed (i.i.d.) Gaussian noise[9]. Current LLR techniques for post-reconstruction improvements utilize an approach where a local patch of size N×N or N×N×N voxels is extracted from each of M time-points. Noise components are suppressed by hard or soft thresholding of the singular values based on the singular value decomposition (SVD) of the N^2^×M or N^3^×M Casorati matrix. In one particular LLR technique, referred to as noise reduction with distribution corrected (NORDIC) principal component analysis (PCA) denoising, the threshold is chosen based on a non-asymptotic random matrix characterization of the thermal noise[16]. Note that for all these LLR denoising techniques, the goodness of such representations depends on the rank of the underlying matrix, where lower rank enhances the likelihood of retaining all relevant image components after thresholding [8].

In this paper, we propose a new transform-domain approach for LLR denoising of MR image series that aims to represent the underlying signal with high fidelity using a lower rank approximation. While a unitary transformation of the images in an MRI series would not affect the global rank, this is not the case for LLR processing. When the images are spatially transformed using a unitary transformation, then the underlying Casorati matrices of the transform domain image series may be represented with lower ranks. Furthermore, unitary transformations ensure that the independent and identically distributed nature of additive Gaussian noise is preserved, thus leaving the random matrix theory framework for parameter-free thresholding unchanged. In this work, we motivate and use this new transform domain approach in the spatial Fourier transform domain of the image series. The lower rank nature of the signal representation in the transform domain is used to enable the denoising of MR image series with as few as 4 volumes, using either complex raw data, complex image data, or magnitude DICOM data and different random-matrix-theory-based approaches to automatically determine the singular value thresholding parameter. Our results show that the proposed transform-domain LLR denoising approach leads to high-quality denoising in MR relaxometry applications with 4 to 12 contrast volumes, improving upon existing image domain based LLR methods[17] that typically require a higher number of volumes in the image series.

## Methods

### Denoising with LLR Models

Let 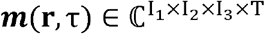 be a complex-valued MR image series, where **r** denotes the location in 3D space, and τ ∈ {1, …, *T* } denotes a particular image in the series across different contrasts or time points. LLR approaches consider *k*_l_ x *k*_2_ x *k*_3_ patches across the image in a sliding window manner. For a given point τ in the image series, this patch is vectorized to y_τ_. These are then concatenated to generate a Casorati matrix Y= [y _1_·,y_τ_·,y_N_ ] ∈ ℂ^M×T^ with *M* = *k*_1_ *k*_2_ *k*_3_ . LLR denoising then aims to recover the corresponding underlying data matrix X, based on the additive noise model

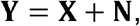

where N **∈** ℂ^M×T^ is Gaussian noise.

LLR models assume that for a given patch across the volume, the data matrix X has a low-rank representation; that is, X can be well-represented, with a low Frobenius norm error, using the sum of a small number of rank-1 matrices.

Based on this assumption, LLR methods perform singular value thresholding using either hard or soft thresholding approaches. Singular values below threshold λ(*j*)< λ_*thr*_ are replaced by λ(*j*) =0. Those above the threshold either remain unchanged (in hard thresholding) or are reduced by λ_*thr*_ (in soft thresholding). For recent works that use random matrix theory (RMT) to characterize the singular value distribution of **Y** or **N**, the threshold λ_*thr*_ is chosen automatically in a parameterless manner based on different criteria, such as matching the tail of a Marchenko-Pastur distribution (as in MPPCA[8]) or ensuring i.i.d. Gaussian noise in **N** by pre-processing and then using the largest expected singular value of a finite Gaussian matrix (as in NORDIC[9]).

While these recent RMT-based LLR denoising methods tackle a major issue in singular value thresholding by allowing parameter-free selection of the threshold without empirical trial-and-error, they still inherently assume that the signal components of the data matrix **X** only have corresponding singular values above the threshold λ_*thr*_, and thus they are assumed to not be conflated with noise and are not removed during processing. This assumption largely relies on the quality of the low-rank representation, which is the focus of this paper.

### Transform domain denoising: A numerical motivating example

As a motivating example consider an image series with 4 volume/images. As a first example, of a full rank series with simple structure, let each image have the same spatial structure (implying no motion), and let each image be constructed from 4 regular partitions with different constant value, and let the relative change from volume to volume be different from partition to partition (Figure 1, row 1). As further examples consider the first example but where the spatial partitioning is unstructured (Figure 1, row 2), and consider a reverse example, where a series of 4 images is created from a quantitative map, using relaxation curves sampled at 4 unique times (Figure 1, row 3).

**Figure 1.**
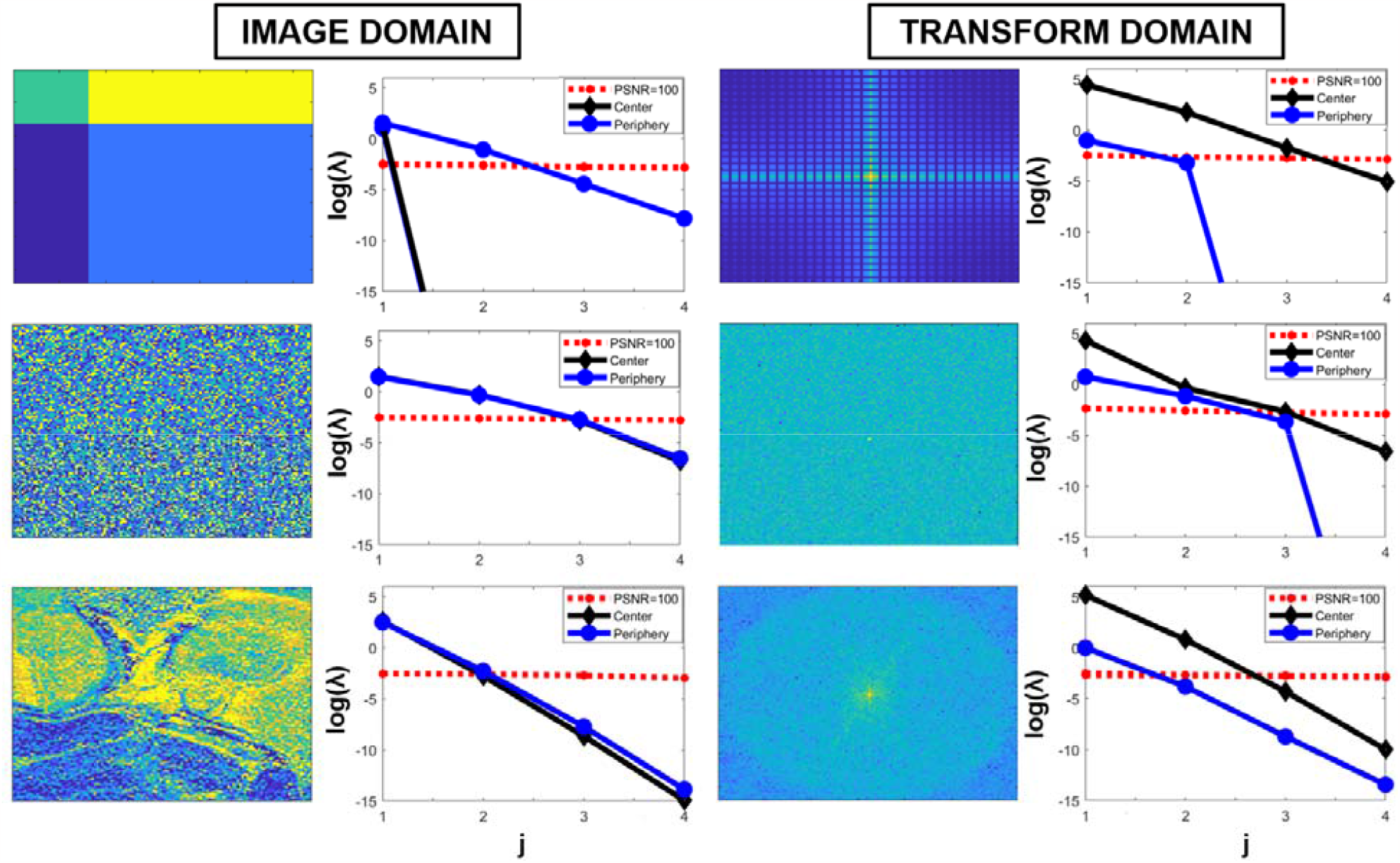
Numerical example of transform domain LLR properties. The left column shows simulated images, the third column shows the Fourier transform of these images, and the second and fourth columns show the singular values for a patch at the center and periphery along with a corresponding noise threshold. The top row is a case of 4 non-interleaved and in time otherwise identical sections, the second row is when the 4 sections are intermixed in a random order, and the third row is when the signal is based on a measured quantitative map. The y-axis for the singular values is plotted logarithmically showing, for the center, an order of magnitude difference between the image domain and the transform domain.

In Figure 1, in addition to the illustrated images in column one, their Fourier transform representations are shown in the third column as an example of a unitary transform. In the second and fourth column, the singular values for the Casorati matrix construction extracted as part of the LLR processing are shown. For reference, the singular values for the Casorati matrix from an i.i.d Gaussian noise image are plotted and scaled relative to an image peak SNR (PSNR) value of 100. For the case of using the LLR denoising techniques directly on the images, the singular values are largely similar to those observed for the center in the transform domain, but are one order lower in magnitude and are thus less distinguishable relative to the noise floor (compared to those in the transform domain). This is in-line with the intuition that the center of the Fourier domain contains most of the contrast information, and LLR characterization in this region corresponds to a global rank constraint on low-resolution images in image domain. For the case of a regular division of homogeneous compartments (top row), in the transform domain, the center is captured by 3 components (with a fourth below the noise floor), whereas for the periphery the signal can be fully represented by 1 to 2 components (the others are orders lower, representing numerical accuracy). For a random division into disjoint compartments (second row), the periphery has more unique components that should be preserved, while for the case of the quantitative map (third row) the periphery, as compared with the center, has more components that will fall below the noise floor and where in the image domain all patches have similar singular values. This example illustrates how, for a case of 4 regular discrete components, the transform space is compressible – largely similar to what would be observed for either of the homogeneous regions (Figure 1, row 1) but not their intersection. For the more conventional applications with structure (Figure 1, row 3), the periphery (high frequency components) also has fewer components to accurately model the underlying signal.

### Transform domain NORDIC (T-NORDIC)

Based on the numerical illustrative example (Figure 1) and the previous results on NORDIC, we propose Transform domain NOise reduction with DIstribution Corrected PCA denoising (T-NORDIC), as illustrated in Figure 2.

**Figure 2.**
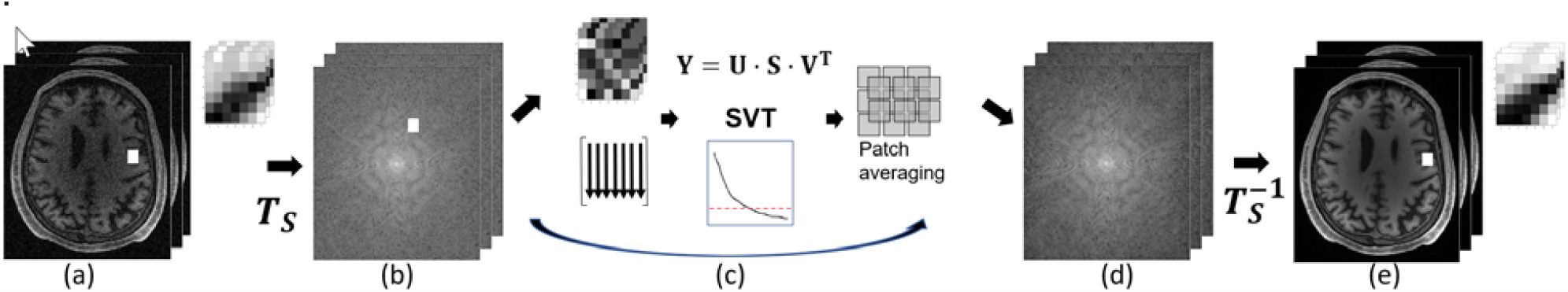
Flow diagram for T-NORDIC. The series of images (a) is processed with a unitary transform ***T***_***s***_. From the transform processed series (b), the same types of transform coefficients are used from each volume (c). For each patch a Cassorati matrix is constructed, SVD is applied, and a hard singular value threshold adapted. Each patch is then averaged together to form a denoised transform domain series (d), and the denoised imaging series is obtained with the inverse unitary transform (e). The zoomed sections in (a) and (e) show the patches used for image-based MPPCA and NORDIC LLR techniques, whereas the zoomed section in (b) show the patches used for T-NORDIC.

**Figure 3.**
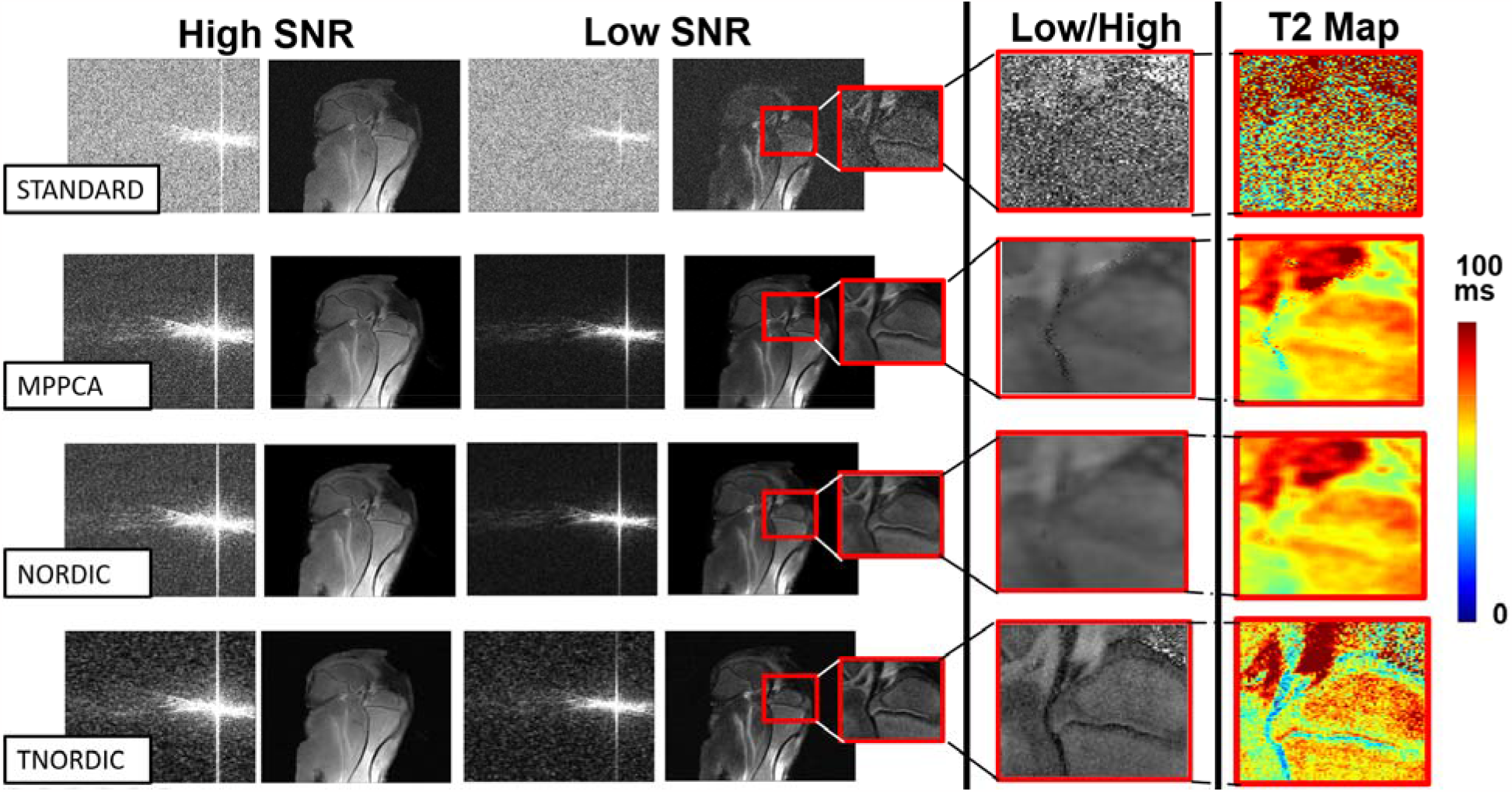
Denoising on single channel data from a MSME T2 map acquisition with 8 echoes and FFT. The first column shows the k-space and image space of the first and last echo, respectively. The second column shows the ratio between the image from the last echo to the first echo. The third column shows the T2 map. The first row shows the acquired data, the second when processed with MPPCA, the third when processed with NORDIC, and the fourth when processed with the proposed T-NORDIC approach.

Following our previous notation, now we consider a spatially transformed version of the image series 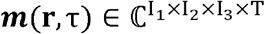, given as

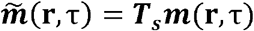

where **T**_S_ is a spatial linear unitary transform, acting on **r** for each point in the image series. Now consider the Casorati matrix 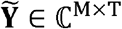 formed by the same process as earlier, but now on 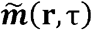. The denoising problem for this Casorati matrix is to recover the transform data matrix 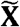 from

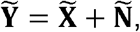

where 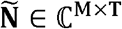 is Gaussian noise. Note that if the noise in m ( r, τ) is i.i.d. Gaussian, so is the noise in 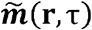, since the transform is unitary, allowing the use of RMT-based selection of the threshold parameter. After singular value thresholding on 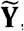, the denoised image patches can be combined to generate an approximation, 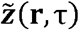, to the spatially transformed MR image series 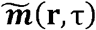. This can then be transformed back to image domain 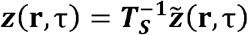, as the final denoised estimate.

For the remainder of this paper, we will consider a spatial discrete Fourier transform (DFT) for **T**_S_. This is a natural choice in MRI as data are acquired in the Fourier domain, and most vendor-supplied filters in MRI are convolutional in nature, and thus implemented as pointwise multiplication in the Fourier domain. Most features related to sampling (e.g., partial Fourier, elliptical, and non-uniform sub-Nyquist sampling) may all be naturally preserved when the DFT is used as a sparsifying transform. While other unitary transforms (such as wavelet transforms) appear equally suited for our purpose, we focus our experiments on the DFT, where the images are processed in the *k*_*x*_*-k*_*y*_*-t* domain instead of the *x-y-t* domain, where *t* broadly refers to the image series dimension. While local patches in the original *x-y-t* domain exploit similarities across image series (e.g., based on an exponential decay for quantitative MRI), the local patches in the *k*_*x*_*-k*_*y*_*-t* domain behave differently. A small patch in the center of this domain captures most of the energy and the underlying contrast information in the image series, which effectively acts like a global singular value decomposition on a very low-resolution image. In contrast, a small patch in outer parts of the *k*_*x*_*-k*_*y*_ domain through *t* captures high-frequency or edge information. For most applications (e.g., fMRI or quantitative MRI), the edge information remains consistent across the image series, thus this representation is able to model the underlying signal with very few components, improving the underlying rank.

### Denoising Experiments

To test the T-NORDIC implementation of transform domain LLR denoising, we compared the denoised results to the standard reconstruction (no denoising), MPPCA (image domain LLR denoising), and NORDIC (image domain LLR denoising) for several relaxation time mapping acquisitions as described below. We considered several different processing cases to highlight the versatility of our approach: (1) single-channel Nyquist-sampled k-space; (2) DICOM images from a Nyquist-sampled acquisition; (3) multi-channel sub-sampled k-space; and (4) publicly available complex-valued multi-channel images.

#### Case #1: T2 mapping using a multi-slice multi-echo (MSME) spin echo sequence (Nyquist -sampled k-space)

First, a fully sampled MSME spin echo T2 relaxation time mapping acquisition was chosen to demonstrate T-NORDIC under Gaussian i.i.d. thermal noise. This acquisition is of a knee joint (i.e., stifle) of a 10-week-old piglet, imaged *ex vivo* at 3T MRI. Sequence parameters were: FOV=120×120 mm^2^; sampling matrix=512×512; in-plane resolution=0.23×0.23 mm^2^; slices=15; slice thickness/gap=2.0/0 mm; TR=1880 ms; TE=13, 26, 39, 52, 65, 78, 91, and 104 ms; partial Fourier=5/8; BW=250 Hz/px; fat saturation; and scan time=10 min.

#### Case #2: T2 mapping using a MSME spin echo sequence (DICOM images)

Second, another a fully-sampled MSME spin echo T2 mapping acquisition was chosen to demonstrate T-NORDIC using the vendor-generated DICOM images. This acquisition is of the knee joint (i.e., stifle) of a four-week-old piglet, imaged *in vivo* at 3T MRI. Sequences parameters were: FOV=128×128 mm^2^; sampling matrix=384×384; in-plane resolution=0.33×0.33 mm^2^; slices=25; slice thickness/gap=2.0/0 mm; TR=4000 ms; TE=11.5, 23.0, 34.5, 46.0, 57.5, 69.0, 80.5, and 92.0 ms; BW=250 Hz/px; fat saturation; and scan-time=16 min.

#### Case #3: T2 mapping using a magnetization-prepared 3D SPACE sequence (sub-sampled multi-channel k-space)

To demonstrate the utility of T-NORDIC for undersampled acquisitions, the 3D SPACE sequence[20] was chosen with its variable flip angle, fast spin echo scheme. This acquisition is of the femoral heads of a 6-week-old piglet, imaged *in vivo* at 3T MRI. Sequence parameters were: FOV=200×100×40 mm^3^; sampling matrix=384×192×40; resolution=0.52×0.52×1.0 mm^3^; TR/TE=2500/217 ms; k-space averaging factor=1.4; partial Fourier; elliptical sampling; BW=501 Hz/px; T2 preparation times=0, 20, 40, and 60 ms; and scan time=11 min.

#### Case #4: T2* multi-echo brain imaging (complex-valued multi-channel images)

To demonstrate the utility of T-NORDIC on anatomical high resolution brain imaging data, a publicly available dataset[21] was chosen. This dataset comprised a 6-echo 3D GRE axial image slab (104 slices) at 0.35mm isotropic (TR. 35ms, TEs: 3.83, 8.2, 12.57, 16.94, 21.31, 25.68, FA 12 degrees, no partial Fourier, 576 x 576 matrix). Whereas the original manuscript combined 4 runs of data (∼1 hour), we selected a single run from the first subject to examine the benefits of T-NORDIC. For both the original data and the data after T-NORDIC, we used the t2smap function from the tedana software[22] to estimate the T2* using a log-linear fit for each voxel in the brain. This T2* estimate was then used to perform a weighted combination of the data to produce a single volume for viewing purposes.

### Implementation details

MPPCA, NORDIC, and T-NORDIC were implemented using a patch size of 5x5x5. This choice was made based on an 11:1 ratio as a good balance between denoising and accurate signal representation[9]. Patch averaging was used with a stride of 2, to ensure at least 8 averages for each denoised value. For MPPCA the threshold selection was implemented on the asymptotic decay of the singular values[8]. For T-NORDIC and NORDIC[9], the threshold selection was implemented as the highest singular value from a Cassorati matrix with i.i.d. entries with a variance estimated from the data[10]. For DICOM data, the noise is estimated from the spectral properties of all Casorati matrices in the data[23].

## Results

### Case #1: MSME spin-echo T2 mapping (Nyquist-sampled k-space)

For an MSME spin-echo acquisition, the impact of the proposed transform domain denoising is shown in Figure 2, where the first (high SNR) and the last echo (low SNR) are displayed. In this case, the SNR reduced on average 90% between the first and the last echo images. The impact on the multi-contrast signal for a single channel fully-sampled k-space is shown for both the image-space and the k-space data. Additionally, the ratio of the low and high SNR images is shown, highlighting the preservation of contrast in T-NORDIC. The effects of edge blurring seen in Figure 2 for MPPCA and NORDIC translates to blurring of the resultant quantitative T2 maps. The ratio of the low SNR to the high SNR image also highlights the challenge of sharing image content between images using a low-rank constraint. The T-NORDIC k-space for the low SNR image illustrates how denoising in the transform domain preserves the magnitude of high spatial resolution features, whereas these are more suppressed with the image space based denoising.

### Case #2: MSME spin-echo T2 mapping (DICOM images)

As a second example of denoising of a MSME spin-echo acquisition, T-NORDIC was applied to standard DICOM data of a piglet knee as shown in Figure 4. The resultant T2 map appears consistently denoised, and from the zoomed insert finer details, consistent with the underlying anatomy, are revealed while subtle heterogenous signal variations are preserved.

**Figure 4.**
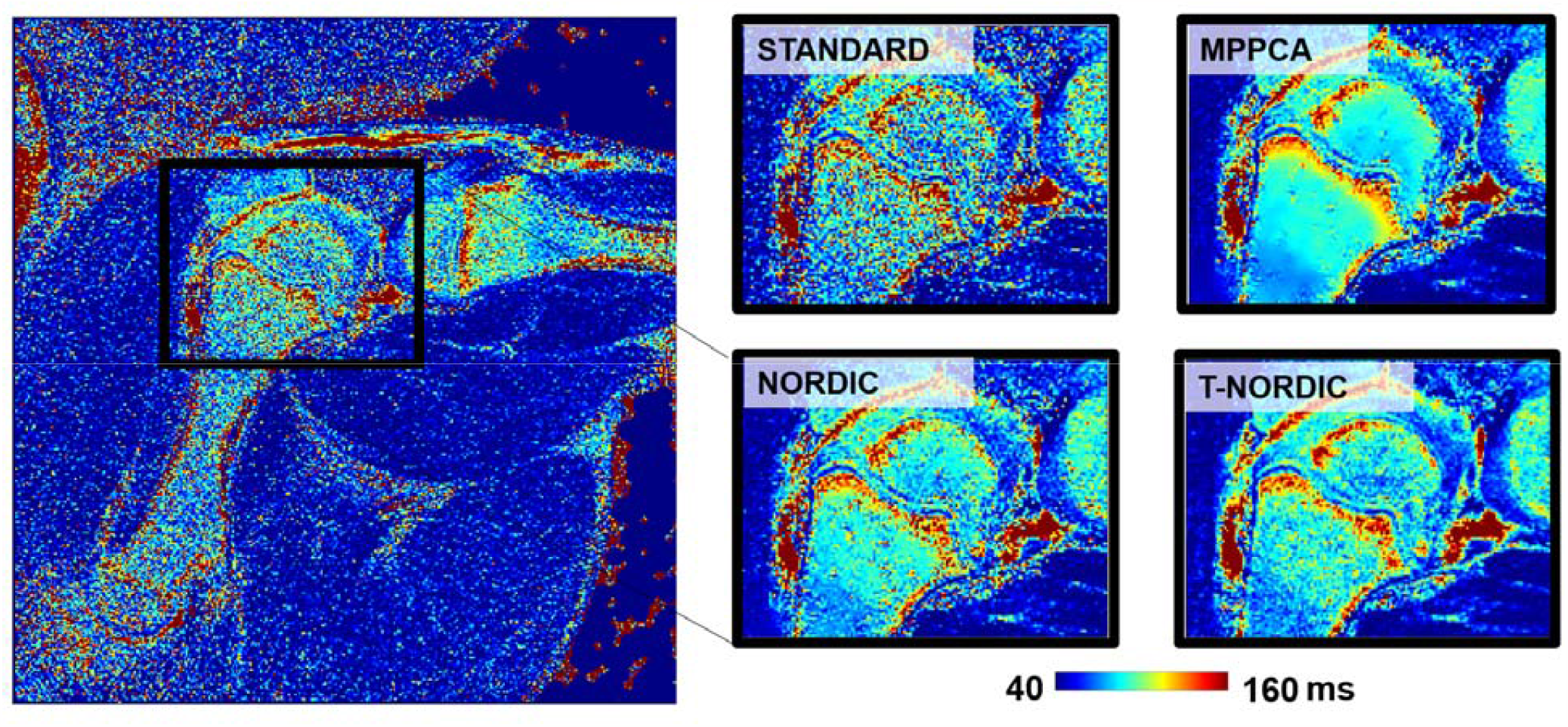
T2 map of a piglet knee generated from DICOM images from an 8-echo MSME spin-echo acquisition. MPPCA, NORDIC, and T-NORDIC were applied to the DICOM images and representative zoomed images of the resultant T2 maps are shown for each of the 4 approaches.

**Figure 5.**
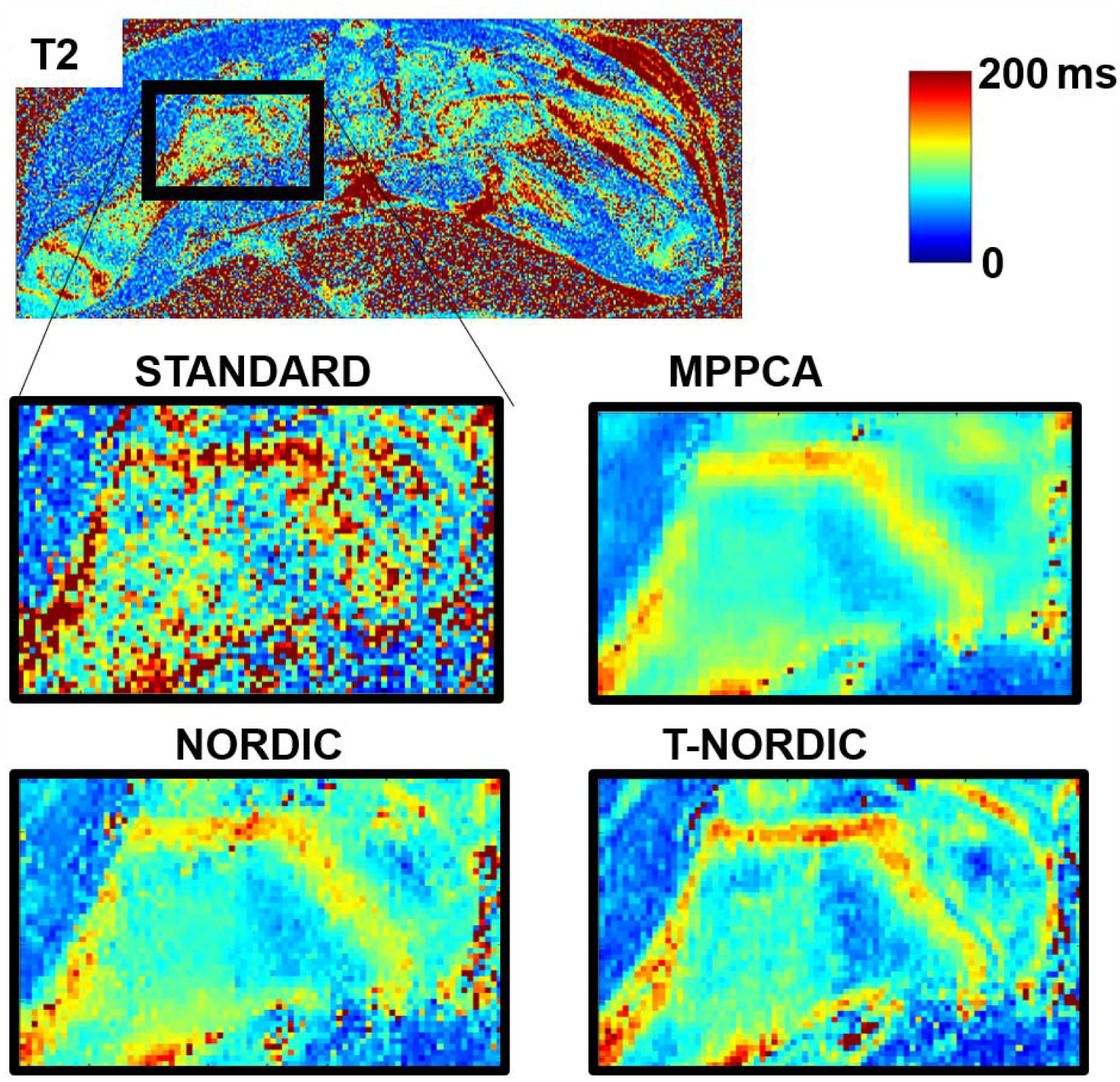
T2 mapping of the hips of piglet with the SPACE sequence and 4 preparation times. The zoomed images show a proximal femur (femoral head and neck) for the standard image reconstruction, MPPCA, NORDIC and T-NORDIC respectively, where the denoising is applied to each of the channels and sampling schemes separately.

### Case #3: Magnetization-prepared 3D SPACE T2 mapping (sub-sampled multi-channel k-space)

To show the impact of denoising on a multi-channel acquisition sequence, Figure 4 shows the effect of denoising before image reconstruction on T2 mapping of a piglet femoral head using a SPACE sequence with T2 preparation. The denoising was applied on the undersampled k-space for each channel independently, and the images were combined with a sensitivity-weighted combination (SENSE-1). The quantitative maps illustrate how all three denoising methods remove noise, but that T-NORDIC most accurately preserves detailed structures and their quantification (such as the relatively long T2 relaxation times of the growth plate and the cartilage overlying the femoral head).

### Case #4: T2* multi-echo brain imaging (complex-valued multi-channel images)

To show the impact of denoising on existing acquired images, the impact of applying T-NORDIC to very high-resolution brain data is shown in Figure 6. As seen in the top row, the estimate of the T2* fit is substantially improved. In the original data, there is a poor fit, with noisy estimates that do not reflect the underlying anatomy (Top Left). This is improved after T-NORDIC (Top Middle), similar to what is obtained by averaging the 4 runs (Top Right). Subsequently, a minimum intensity projection is used over 8 slices to highlight putative venous areas, which can show fine-scale detail. In the original data, this detail is completely lost, with only large veins visible, and overall poor image contrast. After T-NORDIC, however, this detail is recovered and overall image contrast is improved and approaches that of the averaged dataset.

**Figure 6.**
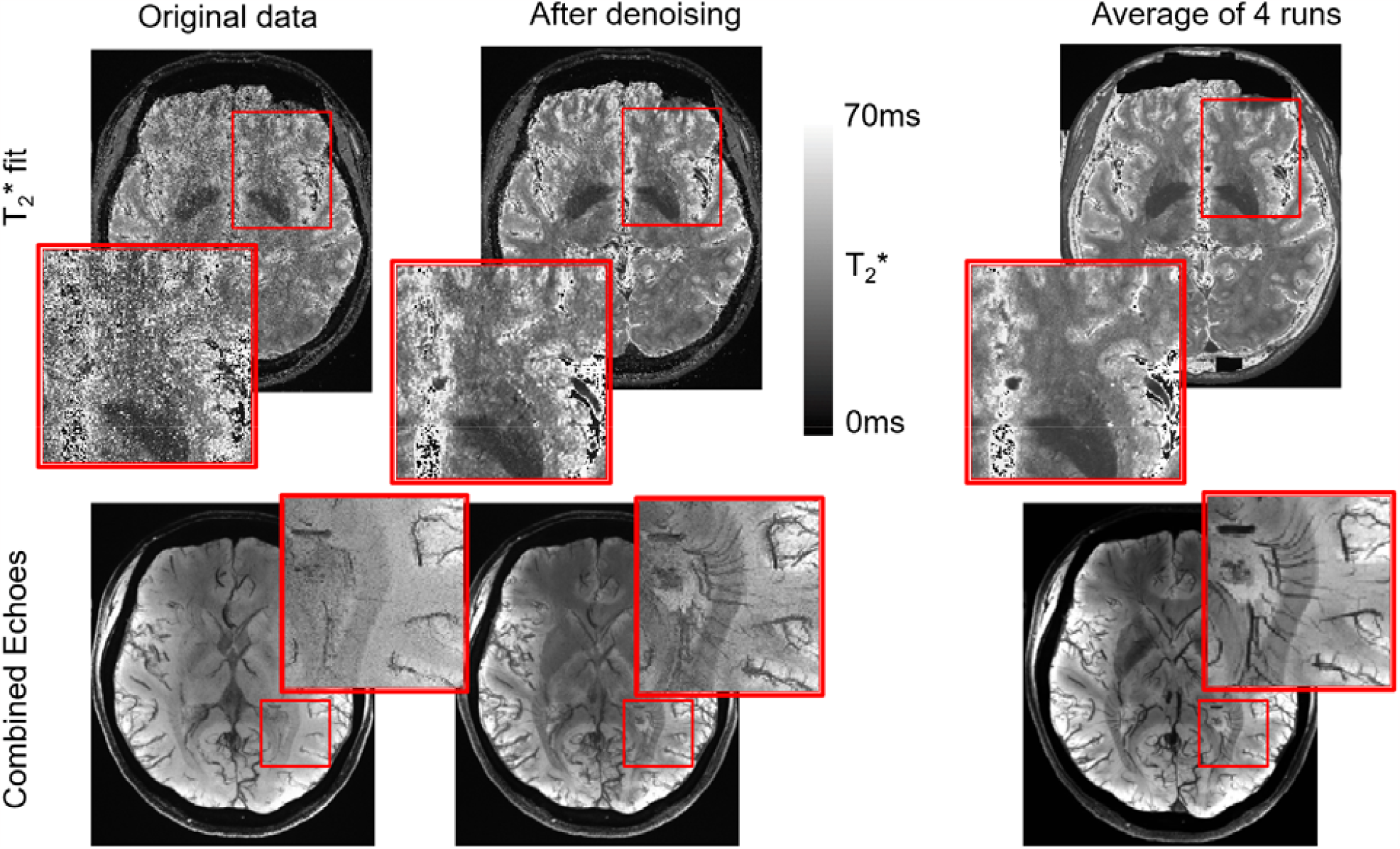
T2* mapping for high resolution brain imaging. Top: Estimate of T2* fit on one run of the original data (Left), on the same data after T-NORDIC (Middle) and on the average of 4 runs (Right). Zoomed in area shows improvement in area with high noise levels. Bottom: A minimum intensity projection of 8 slices of the combined data for the original data (Left), data after T-NORDIC (Middle) and the average of 4 runs (Right). Images show better contrast and zoom-ins show the recovery of fine scale details which match what is seen in the averaged data.

## Discussion

In this study, we proposed the use of spatial unitary transforms for improving the underlying low-rank of Casorati matrices in LLR denoising methods. We further combined this approach with RMT-based parameter-free threshold determination, owing to the preservation of i.i.d. Gaussianity with unitary transforms. Results showed visible improvements over image-domain LLR denoising counterparts in preserving local signal changes when there are large differences in the quantitative values.

The use of LLR techniques have typically required a larger number of images than what is acquired for common parametric mapping protocols. For most parametric protocols, 4 to 8 images are acquired to adequately capture the underlying properties, whereas for most LLR applications, tens of images[17] are needed to adequately preserve the underlying properties. For the MSK applications in areas of more subtle local variability, the transform domain denoising appears to preserve local nuances, whereas the quantitative maps based on image domain methods tend to be locally more homogeneous. For the application of quantitative mapping to low SNR applications, such as high-resolution imaging, the proposed approach will support using fewer averages, thereby reducing scan time and facilitating the integration of quantitative MRI protocols. The ability of the proposed processing to reduce the number of low-rank components for reliable approximation is favorable for quantitative mapping scenarios, and is consistent with the assumption that edges of the image are consistent across frames with varying contrasts.

The theoretical assumption for threshold selection in RMT-based LLR denoising is that the noise is i.i.d. Gaussian, which is consistent with thermal noise in the acquired data. Most MRI images are represented as DICOM images, in which case the noise is either Rician or non-central Chi^2 and may also be spatially varying. The theoretical equations for the thermal noise of these in the transform domain is complicated, yet from the central limit theorem[24] the transform of these distributions may be Gaussian[25]. In such situations, the threshold selection applies and by extension that the denoising in transform domains is more compatible with Gaussian based approaches. Secondarily, since parallel imaging reconstructions, either SENSE[26, 27] or GRAPPA[27], are part of most routine acquisitions, the noise of the interpolated frequencies in k-space is different (normally lower and correlated) from the noise of the acquired data. In such case, the denoising can be performed on the undersampled data before reconstruction, or, if needed, be applied on the post-reconstruction images where the noise in the transform domain should be adequately be accounted for. For GRAPPA based reconstructions, the higher uncorrelated noise of the acquired data is typically preserved and is a better target for noise reduction than the lower correlated noise in the interpolated measurements, but this warrants further investigation

The proposed method has been demonstrated qualitatively for different applications and techniques, for a limited set of images and for an empirically selected set of hyper-parameters. A quantification of the improvement is hard to perform without multiple-averaged data that can be used as a baseline, since measures such as spatial variability that are often used in quantitative MRI applications as a surrogate for SNR improvements inherently favor piecewise smooth denoised outputs, which may also indicate blurring. Future studies with application-specific comparisons to multiple averages are warranted. In this study, we focused on Fourier transform domain processing, and it is worth investigating what other unitary transforms may provide additional benefits. The qualitative investigations were all restricted to a single relaxation mechanism whereas in practice often multiple contrasts, such as T1rho, T2rho are desired or obtained, and the joint integration of these may further compound acquisition time savings. The interaction between the proposed denoising and more complex signal modelling such as DTI or multi-component relaxation[28] also warrants further investigations. We note that both the NORDIC and MPPCA frameworks were introduced for dMRI[7-9, 11, 23] and have been applied to fMRI[10], and in both scenarios the MR image series is sufficiently long to have powerful LLR representations in image domain. Nonetheless, what benefits T-NORDIC over image domain LLR may provide in such scenarios is worth further studies. Some low SNR studies have identified cases where image domain LLR methods lead to biases in functional activation, and the added benefit from T-NORDIC for such scenarios warrants further investigations.

## Conclusion

A transform domain denoising approach that utilizes spatial-temporal properties of noise in a transform domain was proposed to improve denoising quality, especially in scenarios where there are only a few images in the MR image series. Results show that the proposed approach produces high image SNR without deteriorating resolution in a number of quantitative MRI applications, improving upon existing image domain LLR methods. Notably, the proposed approach was applied to different data types, from standard vendor-generated DICOM images to complex multi-channel k-space data, highlighting its potential for widespread use.

## CODE Availability

Code and data to reproduce the figures in this paper will be available at https://github.com/SteenMoeller/ and also via https://www.cmrr.umn.edu/downloads/

## Acknowledgments

The data shown in this paper were collected as part of studies support by the National Institutes of Health (R56AR078315, R56AR078209, P41EB027061, R01HL153146, and R01EB032830). We thank Ferenc Tóth for providing the *in vivo* piglet knee data set.

